# Comparison of progressive hold and progressive response schedules of reinforcement

**DOI:** 10.1101/2022.08.17.504276

**Authors:** Celeste Sofia Alvarez-Sekely, Ana Laura Toscano-Zapien, Paloma Salles-Ize, Maria Almudena Lopez-Guzman, David N. Velázquez-Martinez

## Abstract

Progressive ratio (PR) schedules had been widely used to study motivation to work for a reinforcer. After a post-reinforcer pause, subjects engage pressing a lever until a reinforcer is obtained. However, the discrete nature of lever presses allows alternative behaviors during inter-response time and had lead to the suggestion of several behavioral categories to explain pauses and work time. A progressive hold-down (PH) is incompatible with alternative responses and may allow a precise estimation of work time. Performance of rats trained in both PR and PH that received sucrose or intracranial self-stimulation (ICSS) as reinforcer were compared. We observed that rats mastered the PR and PH schedules. Post-reinforcer pauses, work time and inter-reinforcer time increased as a function of the response or hold requirement. However, rat’s performance suggest that the PH progression may be experienced by the rats as easier that the PR progression. Elimination of consummatory behavior with ICSS reduced PS^R^P and in accordance with predictions of explanatory models of fixed and variable schedules of reinforcement. In the case of PH performance, intermediate requirements leveled off pauses but inceased rapidly on later requirements; since rats controlled pause length and work time was close to hold requirement, time allocation between PR and PH schedules diverged. Finally, the Mathematical Principles of Reinforcement model of Bradshaw and Killeen (Psychopharmacology 2012, 222: 549) rendered a good description of the performance in both PR and PH schedules.

---

Progressive ratio (PR) schedules of discrete response (eg. lever presses) had been widely used to study motivation to work for a reinforcer since the task requires that a subject increases (within a session) the amount of effort (lever presses) expended for subsequent reinforcers. The principal data that many had used is the breaking point (BP), defined as the last completed work requirement before an arbitrary defined time without responding (Hodos 1961; Killeen et al. 2009; Bradshaw and Killeen 2012). After its initial description (Hodos 1961), the effects that several variables have on the PR performance been examined: reinforcement size (Hodos and Kalman 1963; Baron, Mikorski, and Schlund 1992), reinforcer concentration (Bradshaw 2020), effort to depress the lever (Skjoldager, Pierre, and Mittleman 1993), step increase (Hodos and Kalman 1963; Thomas 1974; Stafford and Branch 1998; Covarrubias and Aparicio 2008; Killeen et al. 2009), body weight (Dardano 1973), deprivation level (Ferguson and Paule 1997) and several other variables (Cohen et al. 1994; Ferguson, Holson, and Paule 1994; Rowlett 2000).

The BP has been assumed as the work/effort a subject is willing to pay for a reinforcer, allowing comparisons between the “desirability” of different kind of reinforcers. However, the breakpoint is sensitive not only to interventions that affect the organism’s motivational status but also to those that affect its motor capacity (Skjoldager, Pierre, and Mittleman 1993; Arnold and Roberts 1997; Rowlett 2000), for example, the step increase in the ratio requirement affects the BP as an arithmetic increase leads to larger BPs than a logarithmic progression in the step increase (Killeen et al. 2009). Also, the arbitrary defined time criteria to decide the BP also effects it (Stafford and Branch 1998). In addition, the exclusive use of BP as a performance index of ‘motivation’ put aside many other performance indexes. In particular, the post reinforcement pause has been examined in more detail as an index of performance (Shull 1979; Baron, Mikorski, and Schlund 1992). The Mathematical Principles of Reinforcement (MPR), initially proposed by Killeen (1994) had been suggested as an attempt to overcome the BP as the parameter that resumes the effect of motivational variations and disentangles them from motoric effects in the performance of PR schedules (Bradshaw and Killeen 2012). In their analysis Bradshaw and Killeen (2012) suggested a set of equations where ***a*** (specific activation parameter) expresses the duration of activation induced by the delivery of a single reinforcer and is the primary motivational parameter, being affected by deprivation and incentive motivation; δ (the response time parameter) defines the minimum time that must elapse between the initiation of two successive responses, T_0_ is an initial pause due to post-prandial activity or lassitude, and k is the slope of the linear waiting function. Indeed, there has been several studies that employed the MPR to analyze PR performance (Olarte-Sanchez et al. 2013; Valencia-Torres et al. 2014; Olarte-Sanchez et al. 2015). Here, we describe an attempt to extend the analysis to a Progressive Hold-down schedule (see below).

It has been widely described that emission of discrete responses (eg. pause, response duration, rate) within a run becomes stereotyped; however, there is not guarantee that all the working time WT (time to reinforcer-minus postreinforcement pause (PS^R^P)) is devoted to work. In a seminal attempt to overcome such assumption with discrete responses, Baum introduced the time allocation matching law based on the proportion of time devoted to behavioral alternatives (Baum and Rachlin 1969; Baum 1976; Baum 2013). Based on Baum’s work Shizgal and co-workers introduced the Reward Mountain Model (RMM) (Arvanitogiannis and Shizgal 2008; Hernandez et al. 2010; Trujillo-Pisanty, Conover, and Shizgal 2013; Trujillo-Pisanty et al. 2020) that in a tridimensional space relates opportunity cost, reward strength and work effort using a continuous response that warranties that lever depression corresponds exactly to work effort. Stimulation trains of ICSS have several parameters that determine the total current applied and it has been described that, to a large extent, pulse frequency and amplitude may be a tradeoff to determine the strength of the ICS (Edmonds, Stellar, and Gallistel 1974; Gallistel 1978; Gallistel and Leon 1991; Simmons and Gallistel 1994; Negus and Miller 2014). The counter model has posited that subjects integrates amplitude and frequency as a subjective experience of the magnitude of the stimulation described as reinforcer “strength” (Gallistel, Shizgal, and Yeomans 1981; Gallistel and Leon 1991;

Gallistel et al. 1991; Mark and Gallistel 1993; Simmons and Gallistel 1994). Diverse work with the RMM demonstrated that the amount of work (time allocation) a subject would spend working is related to the joint function of reward strength and opportunity cost (cumulated time or effort to harvest rewards) (Arvanitogiannis and Shizgal 2008; Hernandez et al. 2010; Trujillo-Pisanty et al. 2011; Hernandez et al. 2012; Trujillo-Pisanty, Conover, and Shizgal 2013; Solomon et al. 2015; Solomon, Conover, and Shizgal 2017; Trujillo-Pisanty et al. 2020; Velazquez-Martinez et al. 2022). Previously, there has been attempts to use continuous lever depression as measure of effort in progressive schedules (Progressive Hold-down, PH) where the step increase changes within a session (Bailey et al. 2015; Gulotta and Byrne 2015) or between successive sessions (Rider and Kametani 1984, 1987; Peck and Byrne 2016). As in the case of PR, BPs of PH were also sensitive to both food deprivation and reinforcer quality (Gulotta and Byrne 2015) and reinforcer delay (Peck and Byrne 2016). Hold-duration and distribution has been suggested as an index of motivated behavior (Bailey et al. 2015); also, the PS^R^P has been examined in more detail as an index of performance and was found that its magnitude is related better to the inter-reinforcement time (IS^R^T) than to the WT (Rider and Kametani 1984). Here, we explore whether performance on the PH maintained with sucrose or ICSS may also be described with MPR model of (Bradshaw and Killeen 2012)and if time allocation (the proportion of WT to IS^R^T) PH may also be described by a sigmoid function relating WT to Inter Reinforcement Time (IS^R^T).

## Methods

### 2.1. Subjects

Twenty male Wistar rats (Facultad de Psicología, UNAM), 90 days old and weighing 250-300 g at the start of the study were individually housed in controlled conditions of temperature and humidity, under a normal 12:12 light-dark cycle with light on at 8 am. All rats had initial continuous access to tap water and a pelleted rodent diet (Rodent laboratory Chow 5001, PMI Nutrition International L.LC., Brentwood, USA). At their arrival in the laboratory, their body weights were recorded for 7 consecutive days under free feeding conditions. Throughout the operant sucrose experiment, subjects were maintained at 90% of their initial free-feeding body weights with a correction for their natural growth (in relation to a growth chart from the breeding colony of INNN, Mexico) by adjusting food portion weekly. The rats that received ICSS as reinforcer always had free access to the pelleted rodent diet. Tap water was freely available for all rats in the home cages. All procedures, housing and handling observed the National Institutes of Health guidance for the care and use of Laboratory animals (NIH Publications 8^th^ Ed., 2011) and the study had the approval of the Ethics Committee of the Facultad de Psicologia, UNAM.

### 2.2. Apparatus

The rats were trained in operant conditioning chambers (Lafayette Instruments, Lafayette, IN, USA). One wall of the chamber had a recess into which a peristaltic bomb dispenser could deliver 0.2 ml of a 10% sucrose solution. A retractable lever inserted into the chamber through an aperture situated 8 cm above the floor and 5 cm to the right of the dispenser could be depressed by a force of approximately 0.2 N. A programmable ICSS MED stimulator (PHM-152, MED Associates Inc. Fairfax, VT, USA) provided the train pulses through an electrical swivel (SRO12-0210B10 www.slipringer.com) and a circular orifice on the roof of the chamber and enclosure. The chambers were enclosed in a sound attenuating chest with rotary fans and masking noise. Experimental events and responses were controlled or registered with a MED Associates interface (MED Associates, Inc. Fairfax, VT, USA) and a computer located in the same room.

### 2.3. Procedure :Performance on progressive schedules for sucrose reinforcer

Eleven rats were divided in two groups, one group (response-hold-response, RHRg, N=6) was initially trained on the discrete response PR schedule, then it was trained in the PH schedule and finally, re-evaluated in the PR schedule. The other group (hold-response-hold HRHg, N=5) was trained initially in the PH, then in the PR and re-valuated in the PH.

#### Behavioral training

One week before the start of behavioral training, subjects of both groups were food deprived and their body weight gradually reduced to the 90% of their free-feeding levels. Then, during 2 sessions, they received 0.2 ml of sucrose under a fixed time schedule (FT20 sec), each lever press to either the right or left lever being immediately followed by another 0.2 ml of sucrose. Rats that did not learn to press the lever were shaped manually in two additional sessions. Thereafter subjects were exposed to increasing fixed ratio (FR) schedules (1, 2, 5, 7, 10) for 2 sessions each.

#### PR training

After the last session of FR10 schedule, training in the PR schedules began. The PR schedule was based on an exponential progression: 1, 2, 4, 6, 9, 12, 15, 20,… derived from the formula (5×e^0.2n^)-5, rounded to the nearest integer, where *n* is the position in the sequence of the ratios (Roberts and Richardson 1992; Bradshaw and Killeen 2012). Sessions ended after 90 min or a BP (defined as the last ratio completed before a 10 min period without a response) occurred (Bradshaw and Killeen 2012; Valencia-Torres et al. 2014).Subjects were trained to respond to the right lever until they achieved behavioral stability defined as less than 15% of variability in the BPs during the last 10 sessions. Experimental sessions took place once a day every day (between 12:00 and 16:00), in the light phase of the daily cycle, 5 days per week.

#### PH training

Rats were trained in a cumulative hold-down schedule of reinforcement to press and hold the left lever depressed for increased durations (0.2, 0.5, and 1 s) for 2 sessions each duration. Thereafter, the exponential progression derived from the formula ((5×e^0.2n^)-5)/10 was used: 0.4, 0.6, 0.9, 1.2, 1.5, 2, … On completion of the PH, house light was extinguished, and the session ended. Rats were trained once a day every day (between 10:00 and 16:00), in the light phase of the daily cycle, 5 days per week until they achieved behavioral stability defined as less than 15% of variability in the BPs during the last 10 sessions.

### 2.4. Procedure : Performance on progressive schedules for ICSS reinforcer. Surgery

Rats had anesthetic induction with atropine sulphate (0.05 mg/kg ip) followed 5 min later by ketamine/xylazine (87 and 13 mg/kg ip) and maintained under halothane (Sigma-Aldrich. St Louis, MO, USA)/oxygen vapor mixture (0.5-2% halothane); they were positioned in a stereotaxic frame for bipolar electrode (Plastic One. Roanoke, VA, USA) placement aimed at the MFB at the level of Lateral Hypothalamus (AP: −2.8, ML: ±1.7, DV: -8.9). Electrode was fixed to the skull with dental acrylic; immediately after surgery rats received antibiotic and diclofenac (8 mg/kg ip). At least 1 week was allowed for surgery recovery before operant training.

#### Behavioral training

During the first two days of training rats received increasing intensities of stimulation of 0.1 msec pulses delivered at 100 Hz in a train of 0.5 s duration to identify the highest intensity below the one that produced any sign of discomfort (motor or freezing) to be used as reinforcer. Thereafter, rats were exposed on alternate days to left or right levers with house-light and light above lever turned on during a 30 min session. During such sessions every 20 s a tone of 0.5 s duration was accompanied by intracranial stimulation but any lever press resulted on the immediate delivery of intracraneal self-stimulation (ICSS) simultaneous with the 0.5 s tone; this tone accompanied stimulation and was presented whenever rats obtained a reinforcer. After the ICSS train, responses were ineffective for 0.2 s; during this period and during reinforcer delivery lever lights were turned off. Rats that did not learn to press any lever within 3 days, were shaped manually in additional sessions until they obtained at least 100 reinforcers in a 20 min session. Thereafter subjects were exposed to increasing fixed ratio schedules (1, 2, 5, 7, 10) for 2 sessions each.

#### PR training

After the last session of FR10 schedule, training in the PR schedules began. Training was as described above for sucrose but instead we used ICSS as reinforcer with a waiting time of 6 min to reset PR requirement.

#### PH training

Rats were trained as described for sucrose, but we used ICSS as reinforcer. Rats had a PR and a PH session each day that were separated by 10 min. rats remained in the operant chamber with all lights off and the levers retracted. The start session (PH or PR) alternated randomly each day.

### 2.5. Histology

Rats were anesthetized (sodium pentobarbital; 200 mg/kg IP) and perfused transcardially using saline followed by 4% paraformaldehyde. Brains were then stored at −80 °C. Using a vibratome, 40 μm sections were cut to locate tips of electrode placement, mounted on glass slides and stained with blue methylene. Figure 1 shows tip placements.

**Figure 1.**
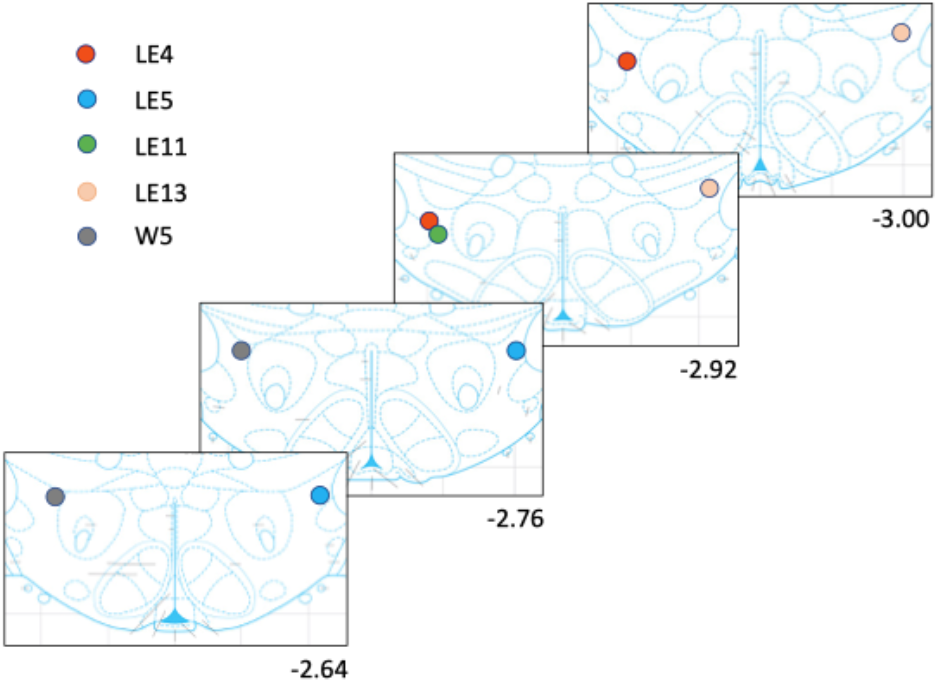
Electrode placement for each rat. Rats had only one implanted electrode either to the right or left side.

### 2.5. Data analysis

To fit the MPR model of Bradshaw and Killeen (2012) we followed the steps described in the appendix of their article and used Excel’s solver (Microsoft 365, v16.56 Redmond, WA, USA). Paired t-test was used to compare pre-to post-conditions; but unpaired t-test was used when comparing between independent groups. Mixed effects Two-way ANOVA (no sphericity assumed, and Geisser-Greenhouse’s correction used) was used to compare between groups at successive steeps of the progressive sequence followed by Šídák’s multiple comparisons test when appropriate. To estimate slope between dependent to schedule requirements we fitted log-log regression lines using the statistical package of PRISM (v9, GraphPad Software, San Diego, CA, USA) was used to fit functions, statical analysis and produce data graphs.

## Results

### PR performance with sucrose reinforcement

There was no difference (paired t_[5]_=0.05874, p= 0.95) on the break points between the pre- and post-conditions of the RHRg, therefore, in the following lines we present the average of both conditions and compare then to the performance of the HRHg. As shown on Figure 2, PR performance was similar in both groups, achieving similar BP (Figure 2A) (unpaired t[9]=0.521, p=0.61) and similar IS^R^T groups was observed for the a, the specific activation parameter (Figure 2C) (unpaired t[9]=2.606, p=0.028); however, no significant differences were observed for δ, the response time parameter (Figure 2D) (unpaired t[9]=0.918, p=0.383), T_0_, the initial pause (Figure 2E) (unpaired t[9]=0.018, p=0.986) or k, the relative readiness to initiate a bout of responding (Figure 2F) (unpaired t[9]=1.344, p=0.212).

**Figure 2.**
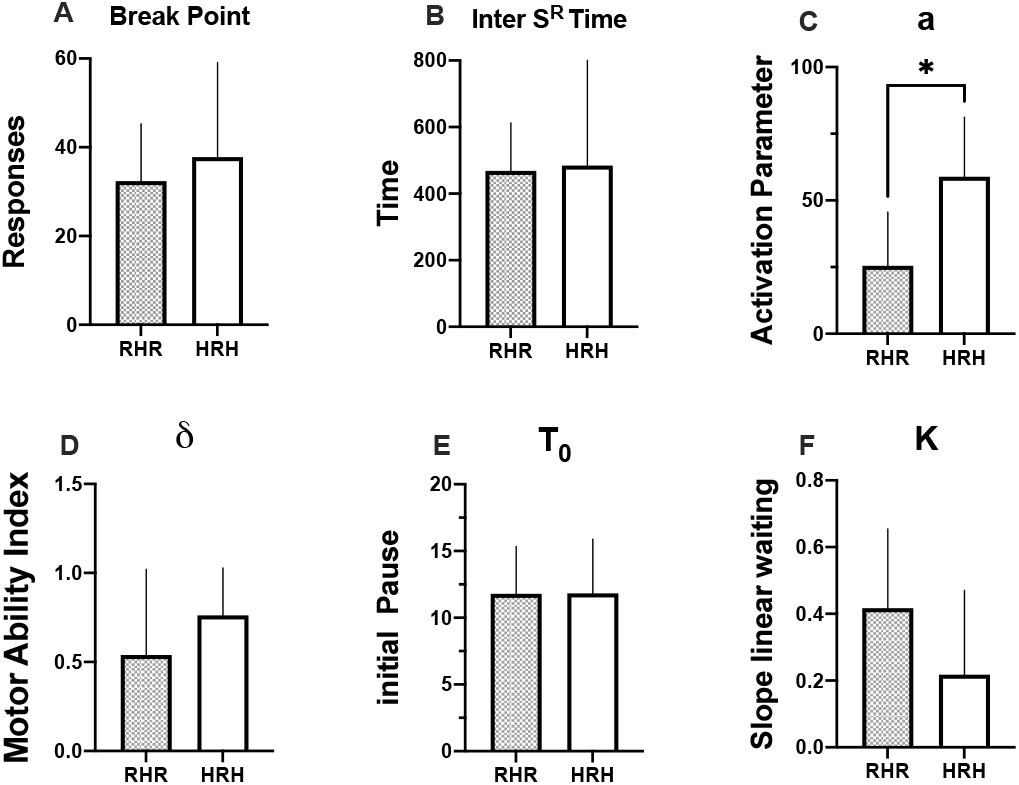
Break points (A), total inter-reinforcement time (B), activation parameter (C), motor ability index (D), initial pause (E) and slope of linear waiting function (F) of the RHR or HRH groups. N= 6 of RHR and N=5 of HRH ± SE.

Figure 3 shows performance measures as a function of ratio requirements; note the ordinate axes are in log scale and that as required ratios increased, number of subjects per group decreased since not all rats completed higher ratios. Figure 3A shows length of PS^R^P on each successive ratio of the PR progression; twoway ANOVA revealed no significant differences between groups (F_[1,9]_=4.849, p=0.055), but significant differences on the step progression (F_[2.180, 16.08]_=6.628, p=0.007), but Šídák’s test revealed differences only for the first pause. When we analyzed WT (Figure 3B) for group differences, two-way ANOVA revealed no significant differences between groups (F_[1,9]_=2.083, p=0.183), but significant differences on the step progression (F_[1.667, 12.00]_=10.08, p=0.004), but Šídák’s test revealed differences only for the sixth requirement.We also compared IS^R^T between groups (Figure 3C); two way ANOVA revealed no significant differences between groups (F_[1,9]_=2.999, p=0.117),but significant differences on the step progression (F_[1.743, 12.86]_=11.72, p=0.002), but Šídák’s test revealed differences only for the first IS^R^T.

**Figure 3.**
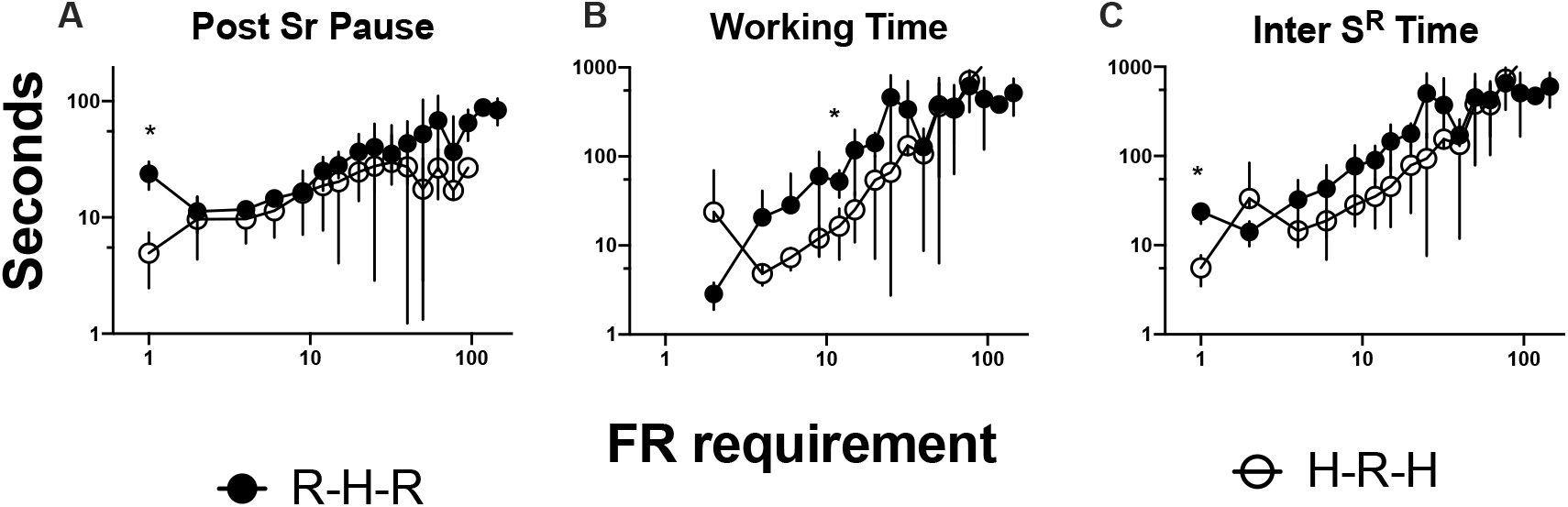
Post reinforcer pause (A), working time (B) and inter-reinforcement time (C) as a function of ratio requirement. N=6 for RHR and N=5 for HRH groups ± SE. Note that as ratio increased, N of groups decreased, and this increases variability.

### PH performance with sucrose reinforcement

Performance measures during PH are presented in Figures 4 and 5. During the PH, time was cumulated for successive holds until the hold requirement was met; therefore, as hold requirement increased it was possible for a rat to press-hold-and-release the lever several times; Figure 3A shows the number of holds emitted during the last requirement completed. Although the group first trained with discrete responses tended to have a larger number of holds-and-releases, no significant differences were found between groups (unpaired t_[9]_=1.894, p=0.091). For the last completed hold requirement, IS^R^T (Figure 3B) was quite similar between both groups (unpaired t_[9]_=0.05944, p=0.954).

**Figure 4.**
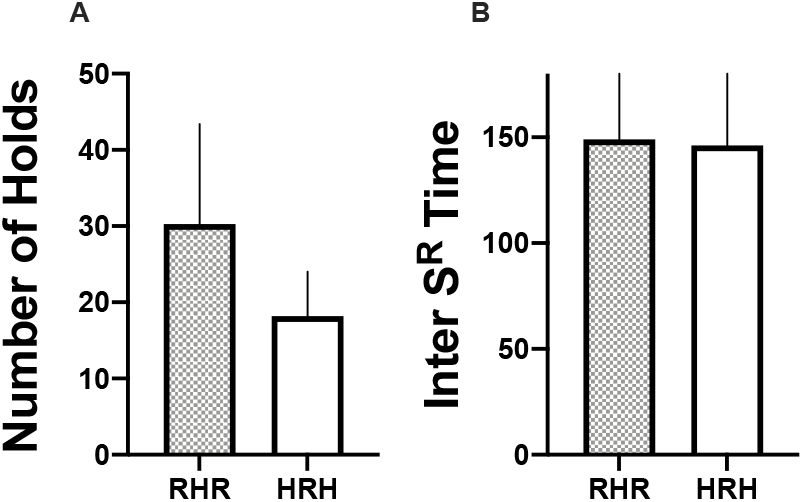
Number of holds (A) and inter reinforcer time (B) during last completed hold requirement. N=6 for RHR and N=5 for HRH groups ± SE.

**Figure 5.**
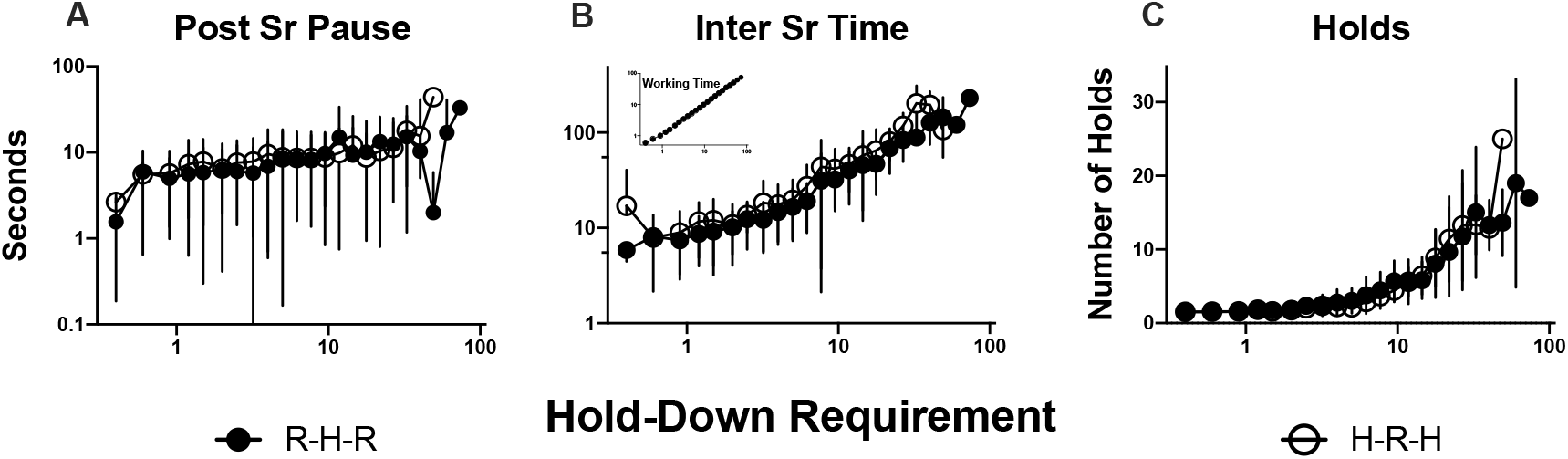
Post reinforcer pause (A), inter-reinforcement time (B), working time (insert B) and number of holds (C) as a function of ratio requirement. N=6 for RHR and N=5 for HRH groups ± SE. Note that as ratio increased, N of groups decreased.

Also, PS^R^P (Figure 5A) was similar between groups and two-way ANOVA confirmed absence of significant differences between groups (F_[1,9]_=0.012, p=0.913), although there were significant differences on the step progression (F_[3.812, 32.05_]=4.153, p=0.008), but Šídák’s test did not revealed differences between groups along the step progression. WT was determined by the completion of hold requirement, so as shown in the insert of Figure 5B it increased orderly as a function of hold requirement. Of more interest is the IS^R^T of the rats; the specific length of the IS^R^T being determined by the hold requirement and the time between successive hold. As shown on Figure 5B two-way ANOVA confirmed absence of significant differences between groups (F_[1,9]_=1.871, p=0.205), although there were significant differences on the step progression (F_[2.588, 21.52]_=27.42, p<0.001), but Šídák’s test did not revealed differences between groups along the step progression. Figure 5C present a similar number of holds between groups; two-way ANOVA confirmed absence of significant differences between groups (F_[1,9]_=0.001, p=0.979), although there were significant differences on the step progression (F_[2.181,18.34]_=31.78, p<0.001), but Šídák’s test did not revealed differences between groups along the step progression.

Since performance under PR and PH were similar between groups, we considered to compare performance under both schedules combining both groups of rats; however, temporal progression produced shorter IS^R^T so, we decided to synchronize both schedules on the basis of the first coincidence of IS^R^T. In Figure 6A notice that the slope of the increase of PS^R^P in relation to the required requirement is larger (mean slope for all data: 0.593; see later for individual slopes) for the PR than for the PH (0.262). Direct observations of the rats indicate that rats keep holding the lever during the initial requirements of the progression while they collect the reinforcer, while none of the rats keep holding the lever during the PR. There was an orderly increase in WT with a slope of 0.983 in the PH, but this only reflects the orderly increase derived from the equation to generate the progression. The progression of the PR also implicated an increase in WT (Figure 6B), but there is no restriction in its slope that was slightly larger (1.070) than for the PH. Notably, the slopes (Figure 6C) of the increase in IS^R^T were similar (PH: 0.8318; PR: 0.928) ; when the slopes were estimated for each rat (Figure 6F), paired t-test indicated no significant (t_[10]_=0.051, p=0.14) difference between schedules with r^2^ slightly superior for PH than PR (Figure 6G), but difference did not reach significance (paired t_[10]_=1.388, p=0.20); however, when we compared the length of the last IS^R^T completed (Figure 6E), clear differences emerged, being the PR IS^R^T significantly larger than that of the PH. Of no surprise, the number of responses in the PR was larger than the number of holds (Figure 6D) and the paired t-test confirmed significant (t_[10]_=3.009, p=0.013) differences between schedules.

**Figure 6.**
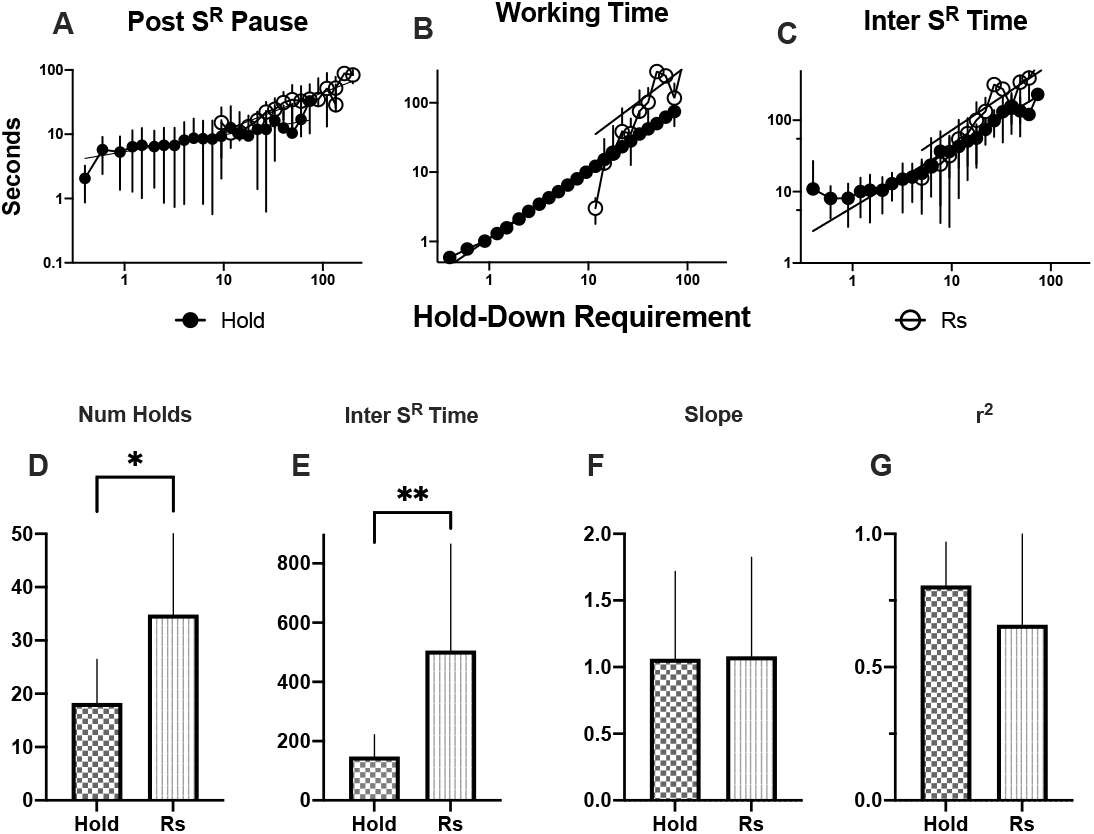
Post reinforcer pause (A), working time (B) and inter-reinforcement time (C) as a function of hold requirement. Abscissas and ordinate in log scale. Number of holds (D) and interreinforcement time (E) of last completed requirement. Slope (F) and r^2^ (G) of log-log regression (see text).N=11 ± SE.

### PR performance with ICSS reinforcement

Figure 7A shows that the PS^R^P under the PR maintained with ICSS starts quite close to 0 but growth faster than in the PR maintained with food. Indeed,two-way ANOVA confirmed significant (F_[3.183,37.60]_=7.899, p<0.001) for the first 3 requirements but not between groups (F_[1,19]_=2.345, p=0.142); such insignificant PS^R^P is related to the non-existent consummatory time related to the ICSS as compared to sucrose reinforcement. Consistent with our interpretation, significant (F_[1.817,20.47]_=15.25, p<0.001) differences in WT (Figure 7B) between requirements but no difference was observed between groups (F_[1,19]_=1.512, p=0.233). Also, significant (F_[1.901,22.46]_=18.16, p<0.001) differences in IS^R^T (Figure 7C) between requirements but no difference was observed between groups (F_[1,19]_=0.984, p=0.334).

**Figure 7.**
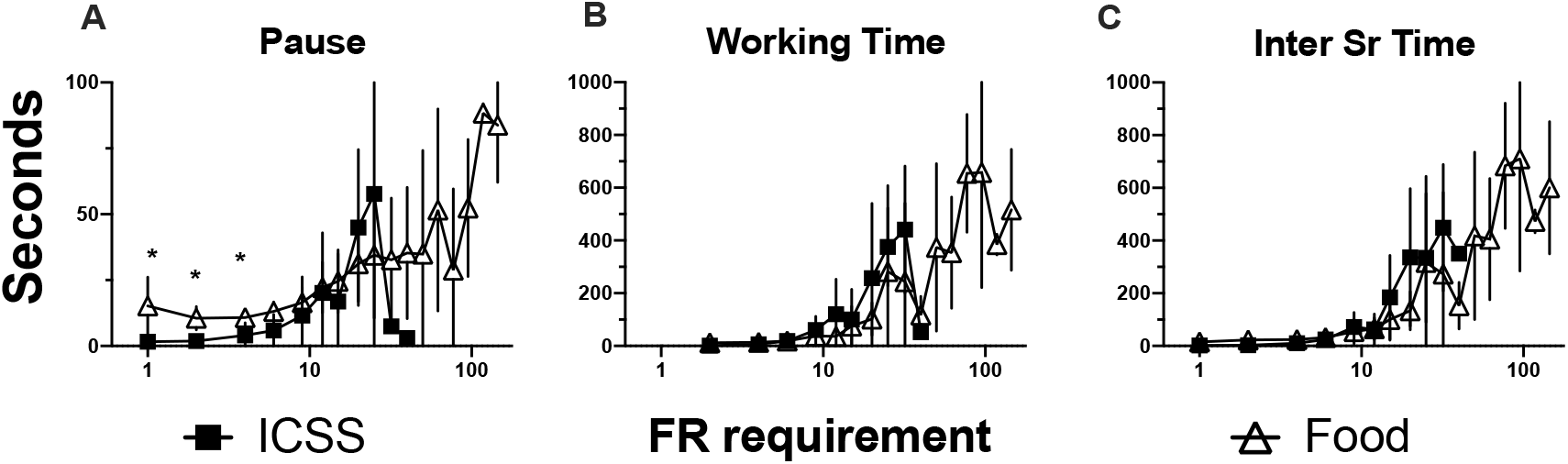
Post reinforcer pause (A), working time (B) and inter-reinforcement time (C) as a function of hold requirement. ICSS N=9, Food N=11 ± SE.

Rats reached significant larger BPs (Figure 8A) with sucrose than with ICSS (unpaired t_[10]_=8.487, p=0.004); however, the IS^R^T during their last completed requirement (Figure 8B) did not reach significance (unpaired t_[19]_=1.362, p=0.192). Consistent with the earlier BP, the motivational parameter (Figure 8C) estimated for sucrose was significantly (unpaired t_[19]_=3.694, p=0.002) larger than for ICSS. However, no significant differences were observed for δ, the response time parameter (Figure 8D) (unpaired t[19]=1-498, p=0.150). Significant differences (unpaired t[19]=8.650, p<0.001) were also observed for T_0_, the initial PS^R^P (Figure 8E) but not for k, the relative readiness to initiate a bout of responding (Figure 8F) (unpaired t[19]=1.639, p=0.118).

**Figure 8.**
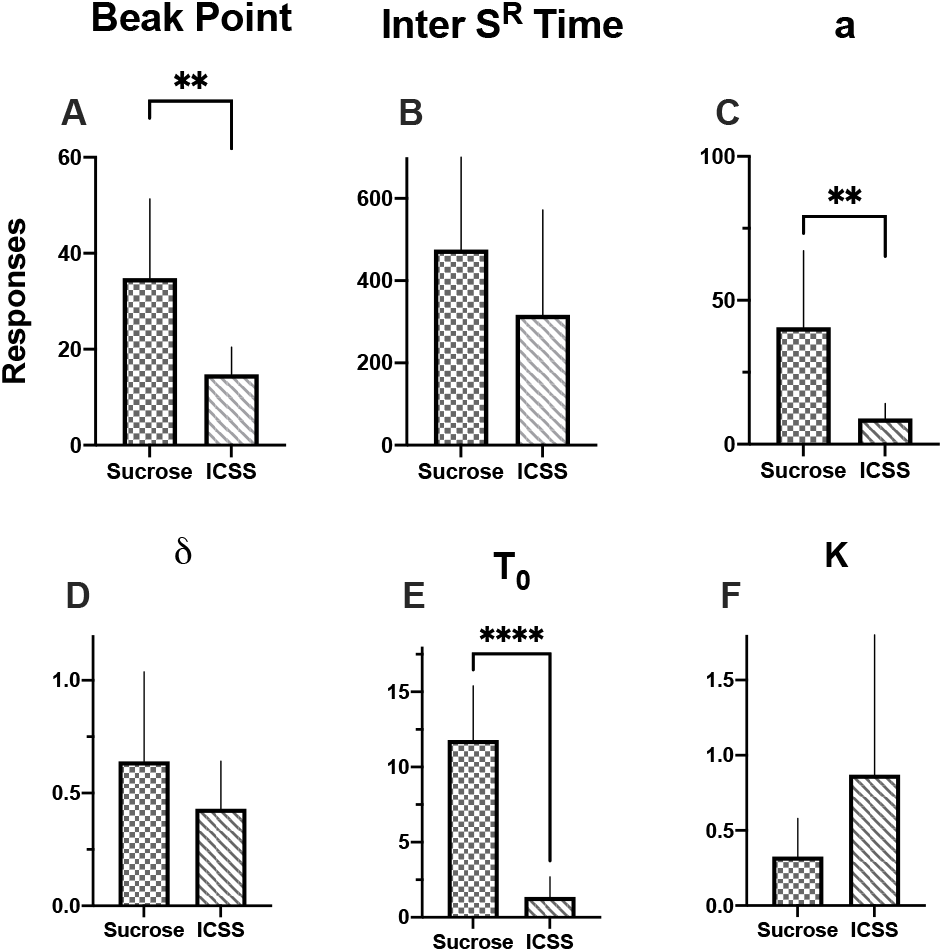
Break points (A), total inter-reinforcement time (B), activation parameter (C), motor ability index (D), initial pause (E) and slope of linear waiting function (F) of the ICSS and sucrose groups. N= 6 of RHR and N=5 of HRH ± SE.

### PH performance with ICSS reinforcement

Figure 9A shows that PS^R^P after ICSS are shorter than sucrose maintained performance and may reflect no consummatory time after ICSS; however, no significant differences emerged after two-way ANOVA between groups (F_[1,18]_=4.042,p=0.06) or step requirements (F_[2.653,35.57]_=2.809, p=0.06).In ICSS-or sucrose-maintained performance, WT was determined by hold requirement as seen on insert of Figure 9B. Of interest is the IS^R^T, the initial IS^R^T being quite short with ICSS, reflecting minimal consummatory time,but later, IS^R^T were quite similar between ICSS and sucrose; two-way ANOVA confirmed no significant differences (F_[1,18]_=2.005, p=0.174) between groups, but significant differences between step requirements (F_[2.474, 33.81]_=30.34, p<0.001); however, we found significant differences (unpaired t_[18]_=0.603, p=0.554) between groups when we compared the IS^R^T of the last requirement completed (Figure 9E). ICSS supported fewer lever presses than sucrose; two-way ANOVA confirmed significant (F_[1,18]_=26.12, p<0.001) differences between groups and significant differences between step requirements (F_[3.217, 44.92]_=16.45, p<0.001); unpaired t-test also confirmed significant (t_[18]_=4.731, p=0.001) differences on the total number of holds of the last requirement completed (Figure 9D). In Figure 9B we plot regression line to group data, but when we fit log-log function to individual data, no significant differences between slopes (unpaired t_[18]_=0.896, p=0.382) or r^2^ (unpaired t_[18]_=0.715, p=0.495) emerged.

**Figure 9.**
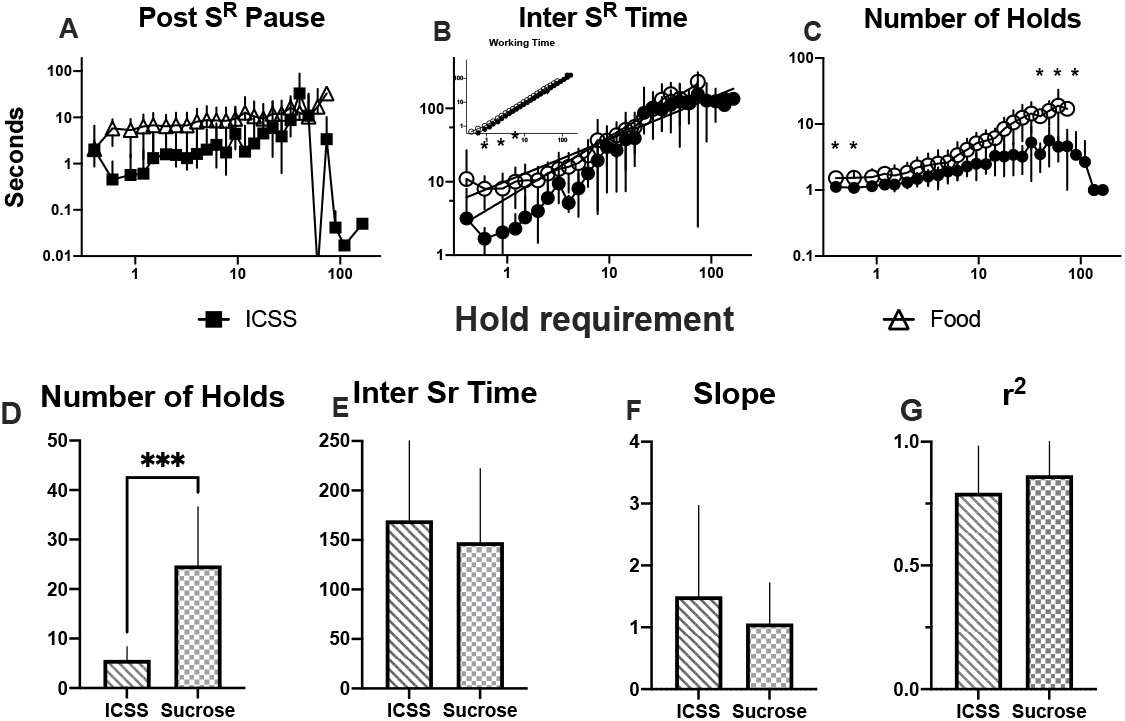
Post reinforcer pause (A), working time (B) and inter-reinforcement time (C) as a function of hold requirement. Abscissas and ordinate in log scale. Number of holds (D) and inter-reinforcement time (E) of last completed requirement. Slope (F) and r^2^ (G) of linear regression (see text). N=9 ICSS, N=11 sucrose ± SE.

When rats were exposed to the initial values of the PH progression, they hold depressed the lever for various step requirements giving very low values of IS^R^T; therefore, to properly compare performance between PR and PH maintained by ICSS, as we did for sucrose, we paired the plots based on equal values of IS^R^T. Pauses (Figure 10A) with PR increased faster than with PH and two-way ANOVA confirmed significant (F_(1,17)_=9.952, p=0.005) differences between programs but not step increments (F_[3.131,26.21]_=2.048, p=0.130). While WT (Figure 10b) had a requirement-dependent increase in PH; WT with PR increased faster than with PH; two-way ANOVA confirmed significant (F_(1,17)_=23.09, p<.001) differences between programs and step increments (F_[2.451,20.97]_=3.315, p=0.047). Also, IS^R^T (Figure 10C) increased faster in PR than in the PH schedule and two-way ANOVA confirmed significant (F_(1,17)_=18.83, p<.001) differences between programs and step increments (F_[3.872,33.70]_=6.084, p=0.001); however, when we fitted log-log function to individual data, no significant differences between slopes (paired t_[8]_=1.598, p=0.074) or r^2^ (paired t_[8]_=1.724, p=0.123) emerged. When we compared the last requirement completed, significant differences for the number of holds or lever presses (Figure 10D: t_[8]_=4.484, p=0.002) but not for IS^R^T (Figure 10E: t_[8]_=1.511, p=0.169) emerged.

**Figure 10.**
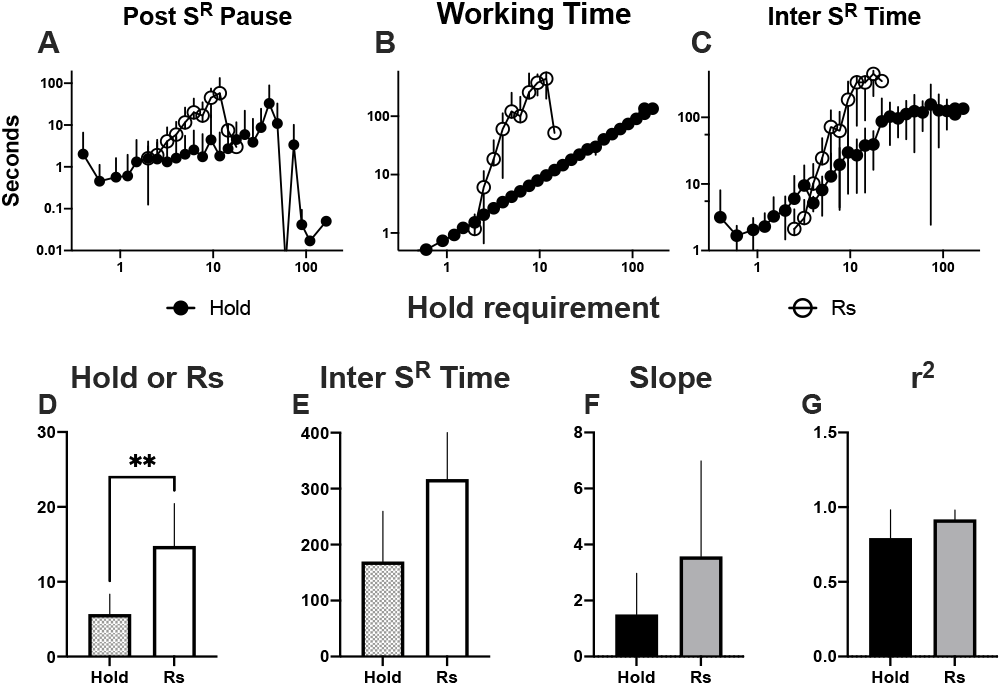
Post reinforcer pause (A), working time (B) and inter-reinforcement time (C) as a function of hold requirement. Abscissas and ordinate in log scale. Number of holds or responses (D) and inter-reinforcement time (E) of last completed requirement. Slope (F) and r^2^ (G) of log-log regression (see text).N=11 ± SE.

As an attempt to use the MPR model to compare performance of PR and PH, we used the reciprocal of the WT and IS^R^T. As the scale used by Bradshaw and Killeen (2012) was minutes we multiplied the reciprocal by 60 and used solver to obtain the parameters presented in Figure 11. The motivational parameter, ***a***, (Figure 11A) estimated for PR-ICSS was lower than for PR-sucrose; however, ***a*** was lower for PH-sucrose and even lower for PH ICSS. Two-way repeated measures ANOVA revealed significant differences on the PR-PH factor (F_[1,17]_=24.34, p=0.001), on the sucrose-ICSS factor (F_[1,17]_=13.08, p=0.002) and on their interaction (F_[1,17]_=10.09 p=0.005); Bonferroni’s test revealed significant differences (p<0.05) between PR- and PH-sucrose and PR-sucrose and PR-ICSS. For ***δ***, the response time parameter (Figure 11B) was similar for PR-sucrose or ICSS; however, it was lower for PH-sucrose and for PH-ICSS gave negative values. Two-way repeated measures ANOVA revealed significant differences for the PR-PH factor (F_[1,17]_=30.43, p<0.001) and for the sucrose-ICSS factor (F_[1,17]_=5.886, p=0.03), but not for their interaction (F_[1,17]_=0.2382, p=0.63); Bonferroni’s test revealed significant differences (p<0.05) between PR- and PH-sucrose and between PR- and PH-ICSS. For ***T_0_***, the initial pause (Figure 11C), the values were similar between PR-ICSS, PH-sucrose and PH-ICSS. Two-way repeated measures ANOVA revealed significant differences on the PR-PH factor (F_[1,17_=47.57, p<0.001), on the sucrose-ICSS factor (F_[1,17]_=97.66, p<0.001) and on their interaction (F_[1,17]_=61.72, p<0.001). Bonferroni’s test revealed significant differences (p<0.05) between PR- and PH-sucrose and PR-sucrose and PR-ICSS. Finally, for K, the relative readiness to initiate a bout of responding, values were similar between PR-ICSS, PH-sucrose and PH-ICSS, and all these were larger than PR-sucrose. Two-way repeated measures ANOVA revealed no significant differences on the PR-PH factor (F_[1,17]_=0.6691, p=0.42), the sucrose-ICSS factor (F_[1,17]_=0.3507, p=0.56) nor their interaction (F_[1,17]_=3.109, p=0.10).

**Figure 11.**
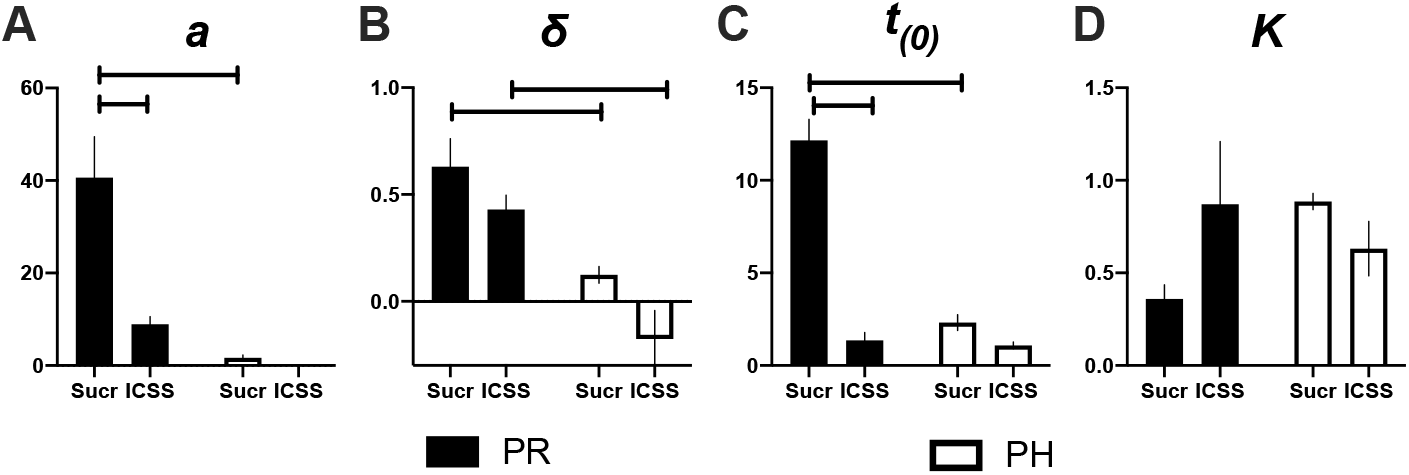
Comparison of motivational parameter (A), motor ability index (B), initial pause (C) and slope of linear waiting function (D) between PR (Rs) and PH (I) groups reinforced with sucrose (S) or ICSS (I). N= 11 for sucrose and N=9 for ICSS ± SE.

Previous work that studied PR performance had shown that PS^R^P is related to WT and IS^R^T (sucrose: Figure 12A, ICSS: 12D) (Rider and Kametani 1984, 1987). Also, it has been shown that the expressing the PS^R^P as a fraction of WT or IS^R^T (Figure 12B, ICSS: 12E) 10934 or a s fraction of the previous ratio requirement (sucrose: Figure 12C, ICSS: 12F) (Roberts and Gharib 2006; Killeen et al. 2009). Some rats achieved larger BPs than others so, as requirement increased, the number of subjects contributing to the means decreased and variability increased; therefore, we fitted the functions to the means that had at least 50% of subjects of each group and the corresponding fitting parameters are in Table 1. We replicate previous findings; worth of notice is that y-intercepts for sucrose are larger than those of ICSS, but in most cases, r^2^ was larger for sucrose than for ICSS. Corresponding plots for PH are presented in Figure 13 and Table 1. As in the case of PR, y-intercepts are larger for sucrose than for ICSS; however, except when expressing the PS^R^P as a fraction of previous IS^R^T, most r^2^ were comparable between sucrose and ICSS

**Figure 12.**
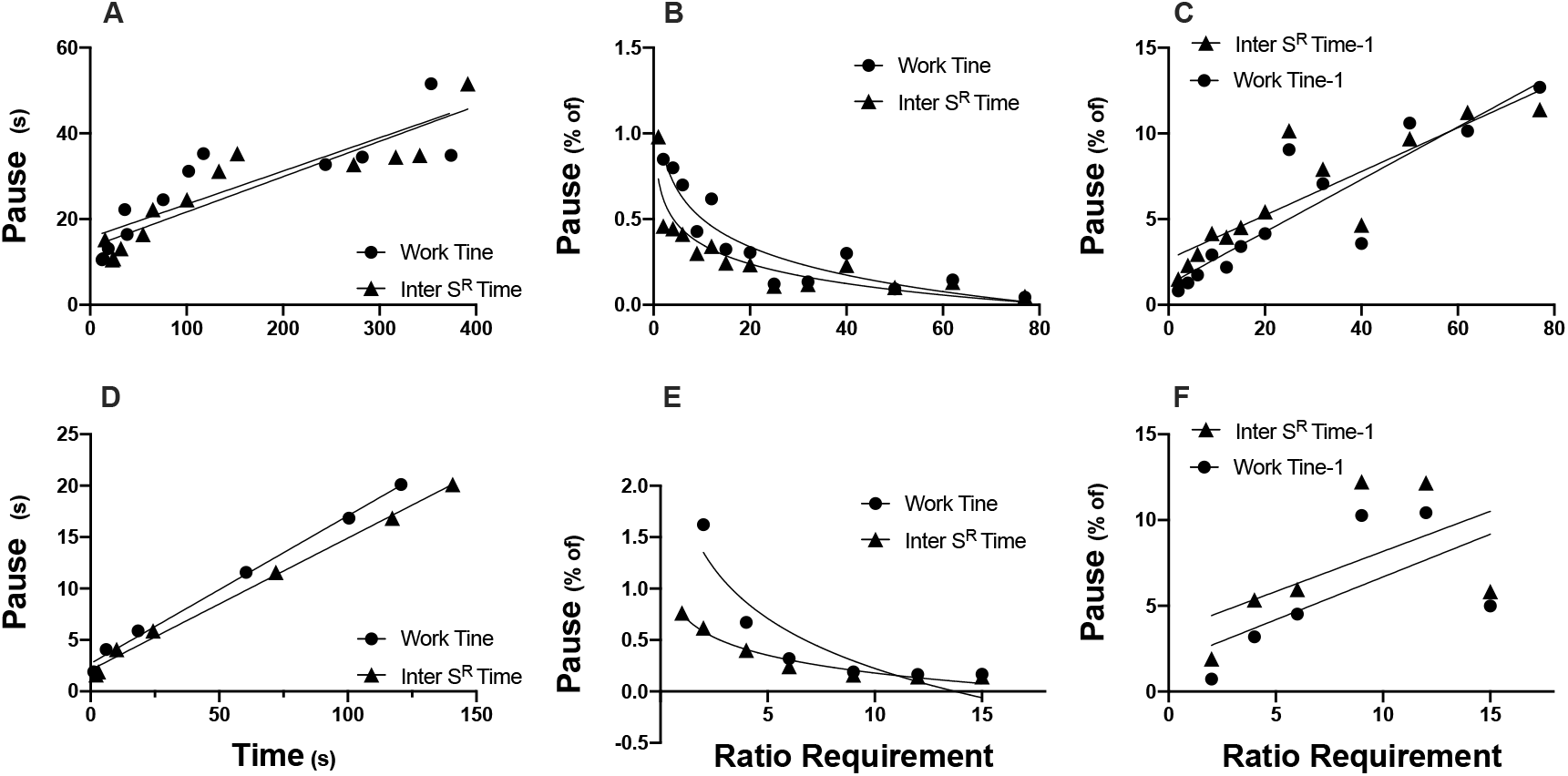
Group mean of sucrose post reinforcer pause in PR as a function of work time (circles) or inter-reinforcement time (triangles) maintained with sucrose (A) or ICSS (D). Pause as a fraction of work time or inter-reinforcement time for sucrose (B) or ICSS (E) or as a fraction of previous requirement work time or inter-reinforcement time for sucrose (C) or ICSS (F) as a function of ratio requirement. See Table 1 for slope, intercept and r^2^.

**Figure 13.**
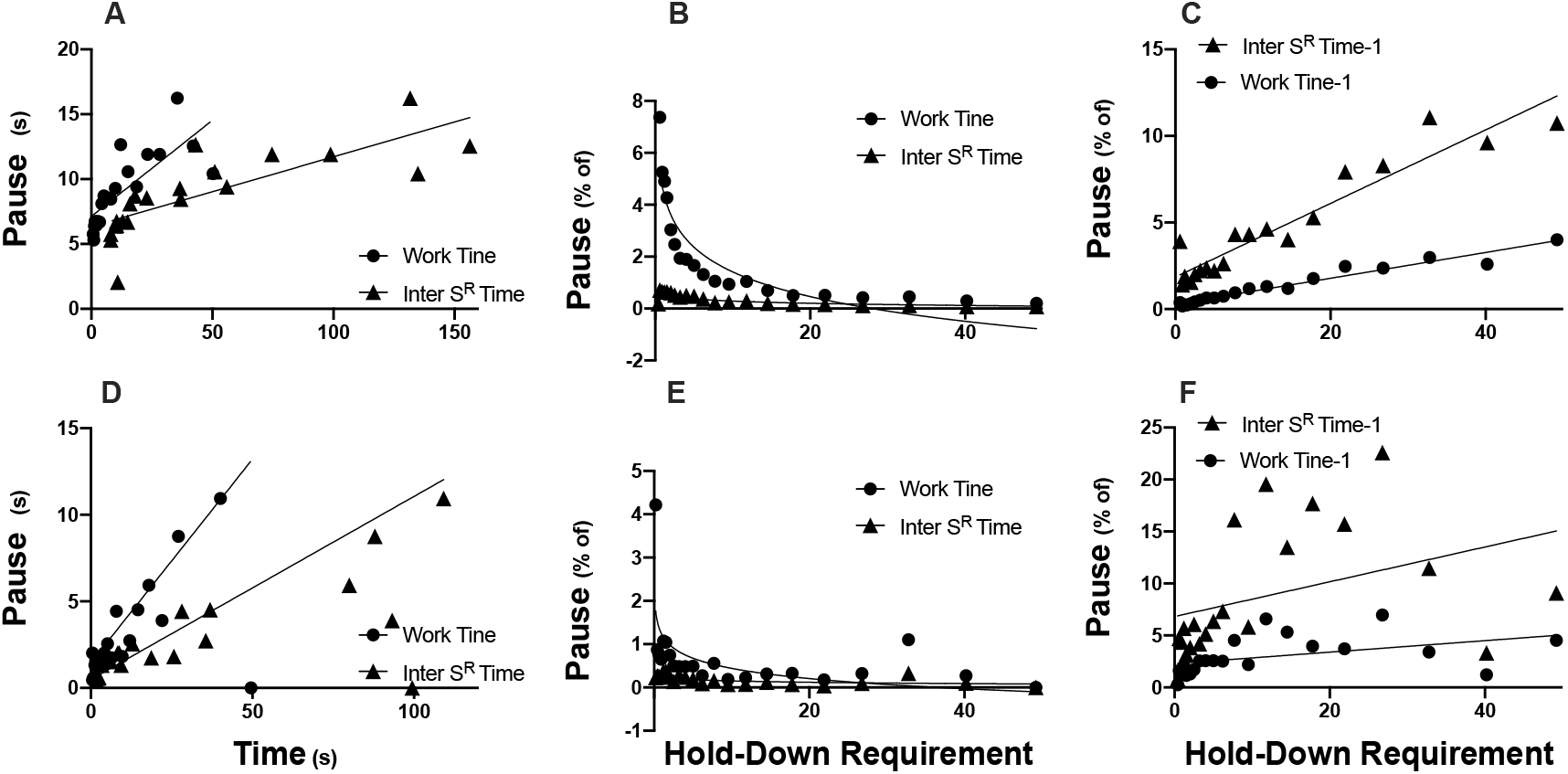
Group mean of ICSS post reinforcer pause in PH as a function of work time (circles) or inter-reinforcement time (triangles) maintained with sucrose (A) or ICSS (D). Pause as a fraction of work time or inter-reinforcement time for sucrose (B) or ICSS (E) or as a fraction of previous requirement work time or inter-reinforcement time for sucrose (C) or ICSS (F) as a function of hold requirement. See Table 1 for slope, intercept and r^2^.

**TABLE 1.** Fitting parameters that correspond to curve fitting of Figure 12 and 13.

|  | Rs |  |  |  | Hold-down |  |  |  |
| --- | --- | --- | --- | --- | --- | --- | --- | --- |
|  | Sucrose |  | ICSS |  | Sucrose |  | ICSS |  |
| Pause | WT | I-S<sup>R</sup>-T | WT | I-S<sup>R</sup>-T | WT | I-S<sup>R</sup>-T | WT | I-S<sup>R</sup>-T |
| Slope | 0.148 | 0.054 | 0.235 | 0.106 | 0.078 | 0.082 | 0.144 | 0.128 |
| Y-intercept | 7.125 | 6.339 | 1.460 | 0.492 | 15.69 | 13.42 | 2.709 | 2.082 |
| $r^2$ | 0.598 | 0.624 | 0.221 | 0.352 | 0.727 | 0.823 | 0.994 | 0.994 |
| Pause as fraction of work time or inter S<sup>R</sup> time |  |  |  |  |  |  |  |  |
| Slope | -3.163 | -0.279 | -0.781 | -0.104 | -0.548 | -0.381 | -1.616 | -0.577 |
| Y-intercept | 4.577 | 0.567 | 1.222 | 0.256 | 1.051 | 0.734 | 1.840 | 0.756 |
| $r^2$ | 0.840 | 0.634 | 0.393 | 0.408 | 0.879 | 0.827 | 0.848 | 0.968 |
| Pause as fraction of previous work time or inter S<sup>R</sup> time |  |  |  |  |  |  |  |  |
| Slope | 0.074 | 0.212 | 0.055 | 0.167 | 0.153 | 0.128 | 0.498 | 0.467 |
| Y-intercept | 0.291 | 1.851 | 2.288 | 6.818 | 1.189 | 2.667 | 1.703 | 3.501 |
| $r^2$ | 0.953 | 0.903 | 0.188 | 0.146 | 0.814 | 0.761 | 0.398 | 0.314 |

We also plot time allocation (TA, proportion of WT to [IS^R^T-reinforcer access]); While TA increased as the response requirement for sucrose increased,there was a small increase of TA as the hold-down requirement for sucrose increased (Figure 14a). With ICSS the increase in TA reached a plateau with the hold requirement, but further increase in hold-down requirement reduced TA (Figure 14b). A similar pattern of TA increase was observed with both glucose or ICSS.

**Figure 14.**
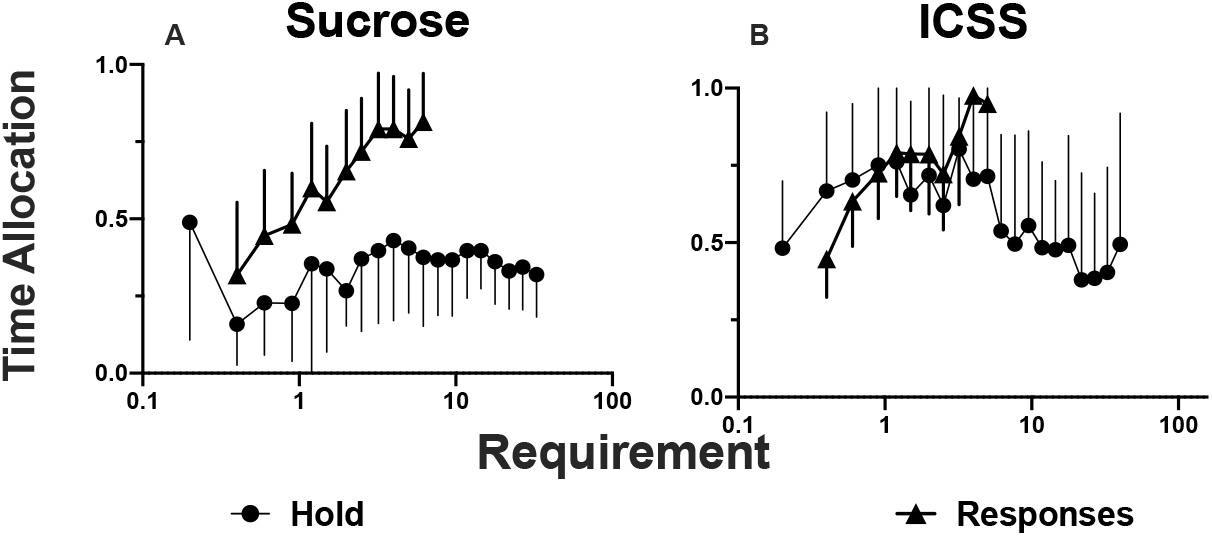
Time allocation (working time as a proportion of available time on each ratio requirement) as a function of schedule requirement. N= 11 for sucrose and N=9 for ICSS ± SE..

## Discussion

We examined the performance of rats under an exponential PR schedule and observed that as the response requirement increased, the PS^R^P, WT and IS^R^T increased when responses were maintained with sucrose, as described previously (Baum 1993; Shull 2004; Killeen et al. 2009; Bradshaw and Killeen 2012). Previous or intermixed PH training sems to have small or no effect on PR performance; with sucrose we used an ABA design for training the PR and PH, but we observed no differences between the first and replicated condition; rats that began their training with PH, showed that the initial PS^R^P tended to be shorter than the PS^R^P on the PR schedule, but on later requirements differences disappeared. So, the PR exerted strong control over responding, and this observation is consistent to those of Cohen et al. (1994) that found that pigeons with a previous history of differential-reinforcement-of-low-rate, variable-ratio or fixed-ratio experience, had shorter PS^R^P than pigeons with only a PR history; however, after prolonged training they found no consistent effects of prior contingencies on responding under the PR schedule.

In general, similar PR performance was observed whether performance was maintained with sucrose or ICSS, the main difference being that the PS^R^P was shorter with the ICSS reinforcer. This difference in PS^R^P may be related to consummatory behavior; rats had to approach, consume, and disengage from sucrose dispenser while they may keep close to the operandum and are ready to respond immediately after ICSS delivery. Some informal observations of rats performing under de PH schedule are consistent with this interpretation; in some cases, when the hold-down requirement was small, rats remained holding down the lever even during the reinforcer delivery.

With the sucrose, rats reached larger BPs than with ICSS. This difference may be related to the time criteria to reset the PR progression; rats that were maintained with sucrose run one session a day and we used 900 s as the time criteria for BP, but those rats that obtained ICSS run 2 sessions a day (PR and PH) and to shorten session time BP time criteria was set to 600 s, and this may contribute to the reduction in BPs. Previously it was described that the number or ratios completed are affected by the BP waiting time criteria, although the BP remained relatively unaffected by the step size (Rider and Kametani 1984).

A lot of work was directed toward the study of PS^R^P on Fixed or Variable Ratio (FR or VR) and Fixed or Variable Interval (FI or VI) schedules of reinforcement. Staddon and Simmelhag (1971) suggested that responding under periodic schedules could be differentiated in two classes of activities: terminal behavior directed toward the scheduled reinforcer and interim behavior possibly maintained by other reinforcers expressed just after reinforcer delivery. Initially, it was suggested that the PS^R^P was mainly composed of interim or nonterminal behavior, while the remining of the IS^R^T or WT was occupied mainly by terminal behavior. However, on a closer examination, it was suggested that an explanation for PS^R^P using discrete responding would require at least three, and possibly four, categories (Shull 1979; Baum 1993; Schlinger, Derenne, and Baron 2008). In PR schedules, initial descriptions showed that PS^R^P increased as the PR requirement increased (Hodos and Kalman 1963); later, a detailed analysis using a PR5 described that PS^R^P increased as a positively accelerated function of the size of the ratio (Baron, Mikorski, and Schlund 1992; Skjoldager, Pierre, and Mittleman 1993). In the case of the PR schedule, we observed that although pauses were larger with sucrose than with ICSS, they keep a similar relation or proportion to WT or IS^R^T and even a similar relation to the preceding ratio requirement (see Figure 12). It was observed that pigeons may engage in self-imposed time-outs during the initial PS^R^P, possibly maintained by other reinforcer(s); but self-imposed time-outs were less frequent during the latter part of a run in PR schedules, where only terminal behavior was observed (Dardano 1973). Baum (1993) showed that in several variable or fixed schedules the PS^R^P covaried directly with the IS^R^T for long IS^R^Ts, but leveled out at an apparent lower limit for short IS^R^Ts; he also observed that curves that related response rate to ratio requirement were apparently bitonic only if the PS^R^P was averaged into the response rate, but If the PS^R^P was excluded, the functions become flat or monotonically increasing; our observations with both sucrose and ICSS (Figures 6, 7, 9 and 10) agree with such observations.

Previous work that examined performance in the PH schedule in some cases used an arithmetic step increment (a constant value) (Rider and Kametani 1984; Gulotta and Byrne 2015), while others used an exponential progression (Bailey et al. 2015). We tested an exponential progression using the same values of step increment than in PR but converted to seconds. In another cases, hold-down was not cumulative, that is, when the rats interrupted the hold-down response before reaching the hold requirement, work time was reset to zero (Bailey et al. 2015; Gulotta and Byrne 2015). Since our purpose was to had the PH schedule similar as possible to the PR, and in the PR effort is cumulative (provided pauses do not exceed the BP criteria), we decided to use a cumulative PH schedule as Rider and Kametani (1984) did. Under such conditions, we observed that the PS^R^Ps in the ICSS PH schedule had larger variability than in the PR schedule (lower r^2^, see Table 1) but they increased faster in relation to WT than in relation to IS^R^T, in accordance with previous findings. Rider and Kametani (1984, 1987) also described that the PS^R^Ps in a non-cumulative PH schedule were relatively short and had a constant duration as the ratio increased, but it increased rapidly near the end of a session for both fixed- and variable-hold requirements and in a further analysis they described that PS^R^Ps duration was positively related to the hold requirement and to the obtained IS^R^T, and to correlate better with the WT, but the intercepts of the lines relating PS^R^P to IS^R^T were consistently closer to zero than the intercepts of the lines relating PS^R^P to hold requirement. In our case, we observed that although the PS^R^P, WT and IS^R^T increased with the hold-down requirement, the increase of the PS^R^P was attenuated in PH compared to the PR schedule, in agreement with previous findings. Also, while IS^R^T progressed at a similar rate for PR and PH with sucrose (Figure 6C) and were similar for PR with sucrose or with ICSS (Figure 9B), its increase was somewhat attenuated in PH maintained with ICSS (Figure 10C). As mentioned above, pauses were shorter with PH than with PR, therefore, we plotted pauses, WT and IS^R^T of PR synchronized to the IS^R^T of PH (see Figures 10); shorter PS^R^P and IS^R^T may suggest that although we programed an exponential step increment in the PH, rats may had experienced such increments closer to an arithmetic than to an exponential step increment. Arithmetic increments produce shallower step increments than exponential increments in PR schedules (Zepeda-Ruiz, Vazquez-Herrera, and Velazquez-Martinez 2020).

Discrete responses may allow the subject to engage in diverse behavioral patterns that may co-vary to determine WT and PS^R^P and, as mentioned above, would require several categories for their explanation (Staddon and Simmelhag 1971;Shull 1979; Baum 1993; Schlinger, Derenne, and Baron 2008). It has been suggested that a more basic unit of measurement than actual numbers of discrete responses may be time allocated to an activity (Baum and Rachlin 1969); then, the hold-down response that excludes any other behavioral responses may truly reflect WT. Hold-down responses had been used successfully by the Reward Mountain Model to have an exact estimation of time allocation and, as commented previously, demonstrated that time allocation is related to the joint function of reward strength (frequency, amplitude or train duration) and the cumulated time or effort to harvest rewards in various electrical-(Arvanitogiannis and Shizgal 2008; Hernandez et al. 2010; Trujillo-Pisanty et al. 2011; Hernandez et al. 2012; Trujillo-Pisanty, Conover, and Shizgal 2013; Solomon et al. 2015; Solomon, Conover, and Shizgal 2017; Trujillo-Pisanty et al. 2020; Velazquez-Martinez et al. 2022) and optical-ICSS (Pallikaras et al. 2022) procedures. Worth to notice is that even the shortest hold requirements rarely were completed with a single lever-holding response, but rather lever was released for very brief periods several times within the IS^R^T (Rider and Kametani 1984). We also observed that hold-down responses were not continuous, but subjects pressed and released the lever several times, but the cumulated number of presses were smaller than in the PR schedule suggesting that inter-response times in PR were larger than the inter-response times under the PH schedule. In some cases, interruptions up to a second had been considered as work time arguing that on such short interruption the rat may be unable to engage in a different activity (Arvanitogiannis and Shizgal 2008; Hernandez et al. 2010; Trujillo-Pisanty et al. 2020). In our case, since we used cumulated lever hold duration we choose not to consider interruptions as continuous holds to determine number of lever releases during execution, but for both PR and PH, our results are consistent with previous observations and predictions of increasing PS^R^P duration as a function of increments of ratio size for both fixed- and variable-ratio schedules and also that the PS^R^P duration may be controlled by the response requirement or the remaining time to reinforcement after the pause (Shull 1979; Mazur and Hyslop 1982; Baum 1993).

In our case, the time allocation (Figure 14) remained stable for intermediate work requirements, but later, it increased as the work requirement increased. In previous work that examined time allocation we observed deceasing time allocation as work requirement increased (e.g. Trujillo-Pisanty et al. 2020; Velazquez-Martinez et al. 2022). However, it should be noted that in previous studies available time for responding (usually enough time to collect a fixed number or reinforcers) was controlled by the schedule, but in the PR schedule, both PS^R^P duration and IS^R^T was controlled by the subject. Therefore, in PR and PH schedules TA adopt a different relation than in those occasion where the subject had a limited time to harvest reinforcers. However, in accordance with the pause analysis presented above PS^R^P duration decreased as schedule requirement increased and allocated time to lever presses or hold-down increased as requirement increased.

As mentioned previously, Bradshaw and Killeen (2012) suggested an analysis for the performance of rats in the a PR schedule; our data with the PR (sucrose or ICSS) fit well with their analysis and the parameters obtained were within the range they described. From our data, it was observed that those rats that first experienced the PH program expressed a larger activation disposition (***a***) than those that began with the PR, but no differences in motor ability (***δ***) or initial pause (***T_0_***) was observed between them. As an approach to analyze PH with the model of Bradshaw and Killeen (2012) and compare it to PR performance we used the reciprocal of time. Rats whose PR performance was maintained with ICSS had lower BPs, and when their performance was compared with the Bradshaw and Killeen (2012)’s analysis: we realize that the activation parameter (***a***) was larger with sucrose than ICSS, but also was larger for PR than PH, there was differences in motor ability (***δ***) between PR and PH schedules but not reinforcer type that may be related to the motor requirements to perform multiple responses or maintain a steady position over the lever. The initial pause (***T_o_***) was minimal with ICSS and PH-schedules, as described above. Finally, worth to notice is that no difference in slope was observed between reward type or schedule used. In the context of the described analysis of PS^R^P, WT and IS^R^T described above, we can conclude that the Bradshaw and Killeen’s analysis renders a good description of the performance in both schedules.

A main limitation of the study is that we used only one value of sucrose concentration or ICSS stimulation. Although it has been shown that ***a***, the motivational parameter, varies with variations of reinforcer magnitude (Bradshaw and Killeen 2012), it may be worth to examine variations of ICSS strength. Also, the first steps in the PH progression and the arithmetic-like slope of WT seems to reveal that rats experienced a low effort performance; therefore, it would be worth to explore a different progression or a progression that omit the early steps of the one that we used in the present experiments.

In conclusion, we observed that rats mastered the PR and PH schedules and their performance in PR schedules was in accordance with previous findings. Performance under the PH schedule was also in accordance with previous findings using PR schedules, but rat’s performance suggest that the PH progression may be experienced by the rats as easier that the PR progression. Elimination of consummatory behavior with ICSS reduced PS^R^P and is in accordance with predictions of explanatory models of fixed and variable schedules of reinforcement. Finally, Bradshaw and Killeen’s analysis renders a good description of the performance in both PR and PH schedules.

## References

Arnold, J.M. and Roberts, D.C.S. 1997. A Critique of Fixed and Progressive Ratio Schedules Used to Examine the Neural Substrates of Drug Reinforcement. Pharmacology Biochemistry and Behavior, 57: 441–447.

Arvanitogiannis, A. and Shizgal, P. 2008. The Reinforcement Mountain: Allocation of Behavior as a Function of the Rate and Intensity of Rewarding Brain Stimulation. Behavioral Neuroscience, 122: 1126–1138.

Bailey, M.R., Jensen, G., Taylor, K., Mezias, C., Williamson, C., Silver, R., Simpson, E.H. and Balsam, P.D. 2015. A novel strategy for dissecting goal-directed action and arousal components of motivated behavior with a progressive hold-down task. Behav Neurosci, 129: 269–80.

Baron, A., Mikorski, J. and Schlund, M. 1992. Reinforcement magnitude and pausing on progressive-ratio schedules. Journal of the experimental analysis of behavior, 58: 377–388.

Baum, W.M. 1976. Time-based and count-based measurement of preference. Journal of the experimental analysis of behavior, 26: 27–35.

Baum, W.M. 1993. Performances on ratio and interval schedules of reinforcement: Data and theory. Journal of the experimental analysis of behavior, 59: 245–264.

Baum, W.M. 2013. What Counts as Behavior? The Molar Multiscale View. The Behavior analyst, 36: 283–293.

Baum, W.M. and Rachlin, H.C. 1969. Choice as time allocation. Journal of the experimental analysis of behavior, 12: 861–874.

Bradshaw, C.M. 2020. Choice between different concentrations of sucrose in an adjusting-magnitude schedule: Evidence for reinforcer-specific value maxima. Behavioural Processes, 181: 104275.

Bradshaw, C.M. and Killeen, P.R. 2012. A theory of behaviour on progressive ratio schedules, with applications in behavioural pharmacology. Psychopharmacology (Berl), 222: 549–64.

Cohen, S.L., Pedersen, J., Kinney, G.G. and Myers, J. 1994. Effects of reinforcement history on responding under progressive-ratio schedules of reinforcement. Journal of the experimental analysis of behavior, 61: 375–387.

Covarrubias, P. and Aparicio, C.F. 2008. Effects of reinforcer quality and step size on rats’ performance under progressive ratio schedules. Behavioural Processes, 78: 246–252.

Dardano, J.F. 1973. Self-imposed timeouts under increasing response requirements. Journal of the experimental analysis of behavior, 19: 269–287.

Edmonds, D.E., Stellar, J.R. and Gallistel, C.R. 1974. Parametric analysis of brain stimulation reward in the rat: II. Temporal summation in the reward system. J Comp Physiol Psychol, 87: 860–9.

Ferguson, S.A., Holson, R.R. and Paule, M.G. 1994. Effects of methylazoxymethanol-induced micrencephaly on temporal response differentiation and progressive ratio responding in rats. Behav.Neural Biol., 62: 81.

Ferguson, S.A. and Paule, M.G. 1997. Progressive ratio performance varies with body weight in rats. Behavioural Processes, 40: 177–182.

Gallistel, C.R. 1978. Self-stimulation in the rat: quantitative characteristics of the reward pathway. J Comp Physiol Psychol, 92: 977–98.

Gallistel, C.R. and Leon, M. 1991. Measuring the subjective magnitude of brain stimulation reward by titration with rate of reward. Behav Neurosci, 105: 913–25.

Gallistel, C.R., Leon, M., Waraczynski, M. and Hanau, M.S. 1991. Effect of current on the maximum possible reward. Behav Neurosci, 105: 901–12.

Gallistel, C.R., Shizgal, P. and Yeomans, J.S. 1981. A portrait of the substrate for self-stimulation. Psychol Rev, 88: 228–73.

Gulotta, K.B. and Byrne, T. 2015. A progressive-duration schedule of reinforcement. Behavioural Processes, 121: 93–97.

Hernandez, G., Breton, Y.A., Conover, K. and Shizgal, P. 2010. At what stage of neural processing does cocaine act to boost pursuit of rewards? PLoS One, 5: e15081.

Hernandez, G., Trujillo-Pisanty, I., Cossette, M.P., Conover, K. and Shizgal, P. 2012. Role of dopamine tone in the pursuit of brain stimulation reward. J Neurosci, 32: 11032–41.

Hodos, W. 1961. Progressive Ratio as a Measure of Reward Strength. Science, 134: 943.

Hodos, W. and Kalman, G. 1963. Effects of increment size and reinforcer volume on progressive ratio performance1. J Exp Anal Behav, 6: 387–92.

Killeen, P.R., Posadas-Sanchez, D., Johansen, E.B. and Thrailkill, E.A. 2009. Progressive Ratio Schedules of Reinforcement. Journal of Experimental Psychology: Animal Behavior Processes, 35: 35–50.

Mark, T.A. and Gallistel, C.R. 1993. Subjective reward magnitude of medial forebrain stimulation as a function of train duration and pulse frequency. Behavioral Neuroscience, 107: 389–401.

Mazur, J.E. and Hyslop, M.E. 1982. Fixed-ratio performance with and without a postreinforcement timeout. J Exp Anal Behav, 38: 143–55.

Negus, S.S. and Miller, L.L. 2014. Intracranial self-stimulation to evaluate abuse potential of drugs. Pharmacol Rev, 66: 869–917.

Olarte-Sanchez, C.M., Valencia-Torres, L., Cassaday, H.J., Bradshaw, C.M. and Szabadi, E. 2013. Effects of SKF-83566 and haloperidol on performance on progressive ratio schedules maintained by sucrose and corn oil reinforcement: quantitative analysis using a new model derived from the Mathematical Principles of Reinforcement (MPR). Psychopharmacology (Berl), 230: 617–630.

Olarte-Sanchez, C.M., Valencia-Torres, L., Cassaday, H.J., Bradshaw, C.M. and Szabadi, E. 2015. Quantitative analysis of performance on a progressive-ratio schedule: effects of reinforcer type, food deprivation and acute treatment with Delta(9)-tetrahydrocannabinol (THC). Behav Processes, 113: 122–31.

Pallikaras, V., Carter, F., Velazquez-Martinez, D.N., Arvanitogiannis, A. and Shizgal, P. 2022. The trade-off between pulse duration and power in optical excitation of midbrain dopamine neurons approximates Bloch’s law. Behavioural Brain Research, 419: 113702.

Peck, S. and Byrne, T. 2016. Demand in rats responding under duration-based schedules of reinforcement. Behavioural Processes, 128: 47–52.

Rider, D.P. and Kametani, N.N. 1984. Interreinforcement time, work time, and the postreinforcement pause. J Exp Anal Behav, 42: 305–19.

Rider, D.P. and Kametani, N.N. 1987. Intermittent reinforcement of a continuous response. J Exp Anal Behav, 47: 81–95.

Roberts, D.C.S. and Richardson, N.R. 1992. Self-Administration of Psychomotor Stimulants Using Progressive Ratio Schedules of Reinforcement. In: A.A. Boulton, G.B. Baker and P.H. Wu (Editors), Animal Models of Drug Addiction, Humana Press, Totowa, NJ.

Roberts, S. and Gharib, A. 2006. Variation of bar-press duration: Where do new responses come from? Behavioural Processes, 72: 215–223.

Rowlett, J.K. 2000. A labor-supply analysis of cocaine self-administration under progressive-ratio schedules: antecedents, methodologies, and perspectives. Psychopharmacology, 153: 1–16.

Schlinger, H.D., Derenne, A. and Baron, A. 2008. What 50 years of research tell us about pausing under ratio schedules of reinforcement. The Behavior Analyst, 31: 39–60.

Shull, R.L. 1979. The postreinforcement pause: Some implications for the correlational law of effect. Advances in analysis of behaviour, 1: 193–221.

Shull, R.L. 2004. Bouts of responding on variable-interval schedules: effects of deprivation level. J Exp Anal Behav, 81: 155–67.

Simmons, J.M. and Gallistel, C.R. 1994. Saturation of subjective reward magnitude as a function of current and pulse frequency. Behavioral Neuroscience, 108: 151–160.

Skjoldager, P., Pierre, P.J. and Mittleman, G. 1993. Reinforcer Magnitude and Progressive Ratio Responding in the Rat: Effects of Increased Effort, Prefeeding, and Extinction. Learning and Motivation, 24: 303–343.

Solomon, R.B., Conover, K. and Shizgal, P. 2017. Valuation of opportunity costs by rats working for rewarding electrical brain stimulation. PLOS ONE, 12: e0182120.

Solomon, R.B., Trujillo-Pisanty, I., Conover, K. and Shizgal, P. 2015. Psychophysical inference of frequency-following fidelity in the neural substrate for brain stimulation reward. Behavioural Brain Research, 292: 327–341.

Staddon, J.E. and Simmelhag, V.L. 1971. The “supersitition” experiment: A reexamination of its implications for the principles of adaptive behavior. Psychological Review, 78: 3–43.

Stafford, D. and Branch, M.N. 1998. Effects of step size and break-point criterion on progressive-ratio performance. Journal of the experimental analysis of behavior, 70: 123–138.

Thomas, J.R. 1974. Changes in progressive-ratio performance under increased pressures of air. Journal of the Experimental Analysis of Behavior, 21: 553–562.

Trujillo-Pisanty, I., Conover, K. and Shizgal, P. 2013. A new view of the effect of dopamine receptor antagonism on operant performance for rewarding brain stimulation in the rat. Psychopharmacology (Berl).

Trujillo-Pisanty, I., Conover, K., Solis, P., Palacios, D. and Shizgal, P. 2020. Dopamine neurons do not constitute an obligatory stage in the final common path for the evaluation and pursuit of brain stimulation reward. PLoS One, 15: e0226722.

Trujillo-Pisanty, I., Hernandez, G., Moreau-Debord, I., Cossette, M.P., Conover, K., Cheer, J.F. and Shizgal, P. 2011. Cannabinoid Receptor Blockade Reduces the Opportunity Cost at Which Rats Maintain Operant Performance for Rewarding Brain Stimulation. The journal of neuroscience, 31: 5426–5435.

Valencia-Torres, L., Bradshaw, C.M., Bouzas, A., Hong, E. and Orduna, V. 2014. Effect of streptozotocin-induced diabetes on performance on a progressive ratio schedule. Psychopharmacology (Berl), 231: 2375–84.

Velazquez-Martinez, D.N., Pacheco-Gomez, B.L., Toscano-Zapien, A.L., Lopez-Guzman, M.A. and Velazquez-Lopez, D. 2022. On the Similarity Between the Reinforcing and the Discriminative Properties of Intracranial Self-Stimulation. Frontiers in Behavioral Neuroscience, 16.

Zepeda-Ruiz, W.A., Vazquez-Herrera, N.V. and Velazquez-Martinez, D.N. 2020. Dissociation between binge eating behavior and incentive motivation. Behav Processes, 181: 104273.

