## Supplementary figures and images for "Comparison of progressive hold and progressive response schedules of reinforcement"

### Fig_1.png

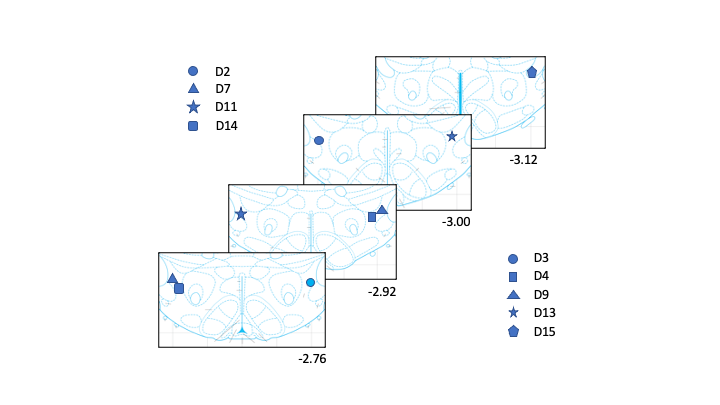
